# Degradation of the intestinal mucus layer by the ETEC protease EatA is species specific determined by the structure of the MUC2 mucin

**DOI:** 10.1101/2025.03.16.643548

**Authors:** Sergio Trillo-Muyo, Brendan Dolan, Frida Svensson, Tim J. Vickers, Liisa Arike, Maria-Jose Garcia-Bonete, Jenny K. Gustafsson, Hans H. Wandell, James M. Fleckenstein, Gunnar C. Hansson, Sjoerd van der Post

## Abstract

Enterotoxigenic *Escherichia coli* (ETEC) infections are a leading cause of diarrheal illness, responsible for an estimated 100,000 deaths annually. ETEC pathogenesis is driven by various virulence factors, including toxins, adhesins, and noncanonical factors such as the protease EatA. The first line of host defense against intestinal pathogenic bacterial infections is the protective intestinal mucus layer. Here, we demonstrate the mechanism by which EatA facilitates access to the epithelial cell surface by degrading the core mucus component MUC2, thereby aiding to the infection. We identify the specific cleavage site region localized at the C-terminal of MUC2. EatA’s protease activity depends on the interaction between two distinct domains, which are uniquely spaced in human MUC2, contributing to species specificity. This was confirmed using a novel chimeric mouse model solely expressing human MUC2, which allowed us to study the role of the mucus layer in the infection of human intestinal pathogens. These findings highlight how ETEC has adapted to specifically degrade the mucus layer of its human host.

## Introduction

Enterotoxigenic *Escherichia coli* (ETEC) infections are one of the major causes of diarrheal illness in low- and middle-income countries ^1^ . While most of the population in endemic regions encounters ETEC, its effects are particularly severe for vulnerable groups. Especially in young children under the age of five where infection can compromise the developing immune system and recurrent episodes may result in malnutrition and stunted growth. ETEC infections are transmitted through contaminated food and water, and subsequently colonize the ileum by adhering to glycolipids and glycoproteins on the epithelial surface to enable efficient toxin delivery. The enterotoxins are responsible for disrupting water and electrolyte balances, leading to diarrhea ^2^. Besides the adhesins and toxins additional colonization factors contribute to ETEC virulence ^3^. These including secreted proteases belonging to the serine protease autotransporters of Enterobacteriaceae (SPATE) family ^4^. SPATEs are prevalent in most pathogenic *E. coli* strains, including enteroaggregative and enterohemorrhagic *E. coli* and are functionally associated with the degradation of host cell receptors and mucus ^5,6^. The main physical barrier in the intestine preventing direct interaction between pathogenic microorganisms and the host epithelium is the mucus layer ^7^. As proximity to the intestinal epithelium is important for adhesion and subsequent toxin delivery a strategy to overcome the mucus layer is a required for infection. We have previously identified that ETEC secretes EatA, a member of the SPATE family that can degrade the MUC2 mucin, the main structural protein component of the intestinal mucus layer ^6,8^. The MUC2 mucin is characterized by its central mucin domains which are composed of multiple repeated sequences composed almost exclusively of proline, serine and threonine (PTS-tandem repeats) that serve as attachment sites for *O*-glycans essential for providing mucus its gel-like properties ^9^. The central mucin domains are flanked by two terminal protein assemblies that are highly stabilized via disulfide bridges and form large oligomeric net-like structures through additional intermolecular disulfide bonds. These distinct features render the protein highly resistant to degradation by members of the commensal microbiota, and only pathogens, including bacteria and parasites have been demonstrated to produce proteases that act on MUC2 and the oligomeric mucus gel ^10–12^. This highlights the necessity of intestinal pathogens to adapt strategies to degrade the mucus to facilitate access to the underlying tissue to increase virulence.

In the present study we provide a detailed mechanistic insight into how the ETEC protease EatA degrades the intestinal mucus layer. Identifying the proteolytic cleavage site and demonstrating the importance of the MUC2 protein structure for substrate recognition, which is dependent on the interaction between multiple domains. Using structural models and a novel transgenic human MUC2 expressing mouse model we demonstrate how variations in protein structure drive species specificity. Providing evidence for the adaptation of pathogenic *E. coli* for degradation of the mucus of its specific host.

## Methods

### Mouse and human subject details

All experimental procedures involving mice were performed in accordance with protocols approved by the laboratory animal ethics committee guidelines at the University of Gothenburg. Mice were housed in a specific pathogen free facility with access to chow and water ad libitum with a 12h light/dark cycle. All mice strains used were on a C57BL/6N background. The Muc2 knockout strain has been previously described ^13^. The MUC2 transgenic mice were generated by homologues recombineering via a pronuclear injection of a BAC clone (RP13-870H17) containing a segment of chromosome 11p15.5 including MUC2 and MUC6. The MUC6 gene was silenced by introducing a TetR cassette ^14^. The mice expressing human MUC2 were crossbred with the Muc2 knockout to generate the MUC2^+/+^/Muc2^-/-^ strain. All animals used for experiments were in the age range between 8 and 12 weeks. Human biopsies were obtained from the distal colon of patients undergoing colonoscopy with no history of inflammatory bowel disease. Only macroscopically normal tissue was included from patients undergoing cancer screening due to occult blood in stool. Approval for the study was granted by the human research ethical committee at the Gothenburg University, and written informed consent was obtained from all study subjects.

### Expression vectors and recombinant protein constructs

The MUC2 N-and C-terminal recombinant proteins were expressed in CHO-K1 cells and purified from spent media as previously described in detail ^15,16^. Expression and design of the truncated MUC2C protein MUC2C-CN is fully described in detail ^17^. The pcDNA3.1(+) plasmid based vector for the MUC2C-N construct was composed of an N-terminal Igk signal sequence followed by a HIS-tag, a Flag-tag site and the MUC2 sequence between amino acid 4256 and 4803 (GenBank: AZL49145.1) with the cysteine residue at position 4786 mutated to an alanine. The mouse Muc2C-N vector was composed similar to the MUC2C-N including the Muc2 sequence between amino acid 3691 and 4248 (UniProt: Q80Z19 v2023_01) with the cysteine residue at position 4231 mutated to an alanine. CHO-S cells cultured in serum free freeStyle CHO growth medium (Gibco) were used for transient protein expression for 72 h. The spent media was concentrated, and buffer exchanged by tangential flow filtration and the protein was subsequently purified by immobilized metal affinity chromatography on 1 mL HisTrap™ HP column (Cytiva) using a Ettan LC system (Amersham Bioscience). The eluted protein was concentrated, and buffer exchanged to 20 mM HEPES, 150 mM NaCl pH 7.4 using 50 kDa molecular weight cut off filters (Vivaspin, Sartorius). The mucin tandem repeats VNTR-PTS1 (x9) and VNTR-PTS2 (x7) were expressed and purified from gene engineered HEK293 cells allowing the control of the *O*-glycans modification, limited to Tn (KO C1GALT1) or STn antigen (KO COSMC/KI ST6GALNAC1) as described in detail ^18^. EatA and the none-catalytic H134R mutant were cloned in a pBAD vector, expressed in *E. coli* and purified from concentrated spent LB media by a ion exchange chromatography followed by size exclusion chromatography as described in detail ^8^. Design and purification of the EatA protein in which the DUF domains spanning amino acid residue 514 to 616 was replaced by eight HIS residues are described in detail previously ^6^. The 34-mer peptide spanning the VNTR-PTS3 between the amino acid 4328 to 4362 of MUC2 was synthesized (JPT) and N-acetylgalactosamine modification were introduced using recombinant GalNAc-Ts transferases 2, 3 and 7 ^19^ .

### Protein gel electrophoresis and Western blot analysis

MUC2C was deglycosylated with β1-3 Galactosidase (P0726, New England Biolabs), α2-3,6,8 Neuraminidase (P0720, New England Biolabs) and *O*-glycosidase (P0733, New England Biolabs) either as a single reaction or sequentially. All reactions were setup according to manufactures instructors without the addition of denaturing buffer and performed overnight at 37°C. For the protease assays recombinant MUC2 was combined with EatA and incubated at 37°C for 1 h on a thermal shaker (300 RPM) if not otherwise stated. The protease activity was quenched by the addition of SDS-PAGE sample buffer (4% SDS, 125 mM Tris-HCl, pH 6.8, 30% (v/v) glycerol, 5% (v/v) bromphenol blue, supplemented with 200 mM DTT for reducing conditions) and heated for 5 min at 95°C. Protein gel electrophoresis was performed on gradient NuPage Bis-Tris mini gels (Invitrogen) with a 4 - 12 or 4 - 15% polyacrylamide gradient and stained by Coomassie (Imperial, Thermo). The protein MW marker used was either PageRuler pre-stained (26619, ThermoFisher), Dual Color (1610374, Biorad) or HiMark pre-stained (LC5699, Invitrogen). Western blot analysis was performed after transfer to PVDF by electroblotting (Turbo transfer, Bio-Rad). Blots were blocked with 5% BSA in PBS-T for 1 h, probed with anti-flag antibody M2 (F3165, Sigma) for 2 h, and detected with anti-mouse IgG1-HRP (4030-05, Santa Cruz). Proteins on blot were visualized by chemiluminescence after the addition of ECL ultra western HRP substrate (Millipore) and the band intensities was analyzed using Image lab (v6.1, Bio-Rad).

### Protein interaction analysis

To determine complex formation the none-catalytic EatA variant H134K or EatA lacking the DUF domain were incubated with MUC2C-N at an equal molar ratio for 1 h at 37 °C. Applied to a Superdex 200 size exclusion column (Cytiva) and eluted isocratically with 20 mM TRIS, 150 mM NaCl pH 7.4 monitoring UV at 215 and 280 nm using an Ettan LC system (Amersham Bioscience). Eluting fractions were analyzed by SDS-PAGE as described. For chemical protein crosslinking analysis EatA was incubated with MUC2C-N or Muc2C-N as above at a concentration of 0.5 µg/µL. Glutaraldehyde solution was added to a final concentration of 0.1, 0.2 or 0.4% and the reaction was terminated after 20 min by the addition of Tris buffer to a final concentration of 50 mM. Cross-linked protein samples were analyzed by SDS-PAGE and visualized by Coomassie (24615, Thermo).

### Mass spectrometry analysis and sample preparations

Trimethoxyphenyl phosphonium (TMPP) protein labeling was performed for the identification of the newly generated MUC2C N-terminal after EatA cleavage. The protease reaction was quenched by the addition of 1 mM phenylmethylsulfonyl fluoride, the protein solution was titrated to pH >8 and <8.5 with Na_2_HPO_4_ and TMPP was added to a final concentration of 100 µM. The reaction was incubated at room temperature for 1 h and subsequently quenched by the addition of 0.1 M hydroxylamine and analyzed by SDS-PAGE. To determine the protein composition and identify TMPP or *O*-glycan modified sites, bands of interest were excised from the gel and destained, proteins were reduced, alkylated, and digested using trypsin (V5111, Promega) overnight at 37°C, as described in detail ^12^. For glycopeptide analysis the protein was first desialylated with a combination of with two broad specific neuramidases (SialEXO, Genovis) and digested into peptides with a Core-1 specific glycoprotease (OpeRATOR, Genovis) in 25 mM ammonium bicarbonate incubated overnight followed by trypsin for 2 h all at 37 °C. The extracted peptides were dried under vacuum and resolved in 0.2% trifluoroacetic acid and analyzed by mass spectrometry. Mass spectrometry analysis were performed by LC/MS/MS using an Easy-nLC (1200, Thermo) coupled to a HF-X orbitrap mass spectrometer (Thermo). Peptides were separated using in-house packed columns (150 x 0.0075 mm) packed with Reprosil-Pur C18-AQ 3 μm particles (Dr. Maisch). Peptide were separated using a 5 to 35% gradient (A 0.1% formic acid, B 0.1% formic acid, 80% acetonitrile) in 30 minutes. Full mass spectra were acquired over a mass range of minimum 350 *m/z* and maximum 1600 *m/z*, with a resolution of at least 60,000 at 200 *m/z*. The 12 most intense peaks with a charge state ≥2 - 5 were fragmented by HCD with a normalized collision energy of 27%, and tandem MS was acquired at a resolution of 17,500 and subsequent excluded for selection for 20 seconds. Peaklists were generated from raw mass spectrometry data using MSConvert and searched using Mascot (v2.2.04, Matrix Science) for peptide identification and Byonic (v2.10.21, Protein Metrics) for glycopeptide identification. Mascot was used with the following settings for standard protein identification fixed modification carbamidomethylation (C) and variable modification oxidation (M), enzyme trypsin with a maximum number of two miss cleavage and mass tolerance was set to 10 ppm for precursor mass and 20 ppm for fragment ions. For newly generated N-termini identification the search settings were adapted to include TMPP (N-terminal, Y and K) and semi trypsin for enzyme specificity. All peptide searches were performed towards an in-house protein database including the recombinant constructs and EatA (<u>Mucin DB</u> <u>v2</u>) and only peptides with a score >30 were considered. Byonic parameters were set to semi specific for C-terminal cleavage trypsin (KR) and N-terminal OpeRATOR (ST) with a maximum number of miss cleavage up to two, mass tolerance of 10 ppm for precursor mass and 10 ppm for fragment ions. The searches we performed against the mucinDB and a list of 70 common human *O*-glycans was selected for considered glycan modifications and only glycopeptides with a score >300 were considered. The unmodified or GalNAc modified 34-mer PTS3 peptide (JPT) was resolved in PBS and further diluted with 0.2% trifluoracetic acid to 2 µM for analysis by LC-MS/MS using the same instrument settings as described. Protease cleavage was validated by combining 100 µM PTS3 with up to 1 µg EatA, incubated at 37 °C overnight till up to 24 h. The reaction was quenched by the addition of 0.2% trifluoracetic acid and analyzed by LC-MS/MS.

### Ex vivo analysis of EatA mucus degradation

*Ex vivo* analysis of EatA was essentially performed as described previously ^20^. Following euthanasia, mouse distal colon was dissected and collected into ice-cold oxygenated (95% O_2_ and 5% CO_2_) Kreb’s buffer (116 mM NaCl, 1.3 mM CaCl_2_, 3.6 mM KCl, 1.4 mM K_2_HPO_4_, 23 mM HCO_3_^-^ and 1.2 mM MgSO_4_). Tissue was opened along the mesenteric border, and after removal of the fecal material and the longitudinal muscle tissue was mounted in a horizontal perfusion chamber ^21^. Human sigmoid colon biopsies were collected into Kreb’s transport buffer and mounted in a horizontal perfusion chamber as for mouse tissue. Tissue was maintained at 37°C throughout the duration of each experiment and was perfused basolaterally with Kreb’s glucose buffer (Kreb’s buffer + 10 mM D-glucose) with Kreb’s mannitol buffer (Kreb’s buffer + 10 mM D-mannitol) added to the apical surface. Both human and mouse tissue were counterstained with Syto9 (10 µM; ThermoFisher) for 5 minutes and the mucus layer was visualized by the addition of 1 µm FluoSpheres™ carboxylate-modified microspheres (625/645; ThermoFisher). 10 ng/ml of EatA was added to the apical Kreb’s mannitol buffer and the integrity of the mucus layer was monitored on an upright LSM900 confocal microscope (Carl Zeiss) using a Pan-Apochromat x20/1.0 DIC 75 mm lens (Carl Zeiss). Following the acquisition of an initial scan before addition of the protease, z-stacks were acquired every 3-5 minutes for up to 45 minutes using Zen Blue software (version 3.1; Carl Zeiss). Control analyses were performed on paired mouse or human tissue treated with the non-catalytic EatA variant H134K. To quantify mucus integrity, beads and tissue surfaces were mapped to isosurfaces using Imaris (version 9.5; Bitplane) as described previously ^22^. Data regarding the bead position in relation to the tissue surface over time was then extracted and analyzed to generate mucus thickness and normalised bead positional data (Prism version 10.3.1; Graphpad).

### In-silico docking and molecular dynamics

The EatA DUF domain structure was predicted using AlphaFold2 ^23^ . The cryoEM structure of MUC2 VWCN-D4-VWC1 (PDB: 7QCN) was modified with Modloop ^24^ in order to complete the missing loops. The glycosylation sites were modified ensuring that all of them were occupied with two sugar molecules, two N-Acetylglucosamines (GlcNAcβ1-4GlcNAc) in *N*-glycosylation sites and galactose and N-Acetylgalactosamine (Core 1: Galβ1-3GalNAc) in *O*-glycosylation sites. The simplified glycans circumvented the risk of masking interaction surfaces due to their high flexibility not being considered by the docking programs, while blocked the regions where they are directly attached. The *in-silico* docking was performed based on shape complementary using PatchDock ^25^ and further refined and ranked with FireDock ^26^ . The full-length EatA structure also predicted by AlphaFold2 and a model of MUC2 from the VWCN domain to CK were aligned to the docking models in order to check their feasibility ^17^ . The DUF domain was replaced by the full length EatA in the best ranked model. The small clashes were solved by a short molecular dynamic simulation (MD). To carry this it out, the complex was prepared for MD using the Glycan Reader and Modeler tool from CHARMM-GUI ^27^ . It was solvated with the TIP3P water model in a cubic simulation box under periodic boundary conditions, neutralized and adjusted to 150 mM NaCl. The generated CHARMM36m force field parameters were used in GROMACS 2022.4 ^28^ to perform the energy minimization, equilibration, and production simulations at 303.15 K. The production was simulated for 5 ns. The PTS3 region of MUC2 chain B was modeled in Coot ^29^ extending it from the VWCN N-terminal residue towards the EatA active site. The PTS was glycosylated using GLYCAM and the resulting model was manually curated in order to remove ring piercing artifacts. The lateral chains of affected residues were rebuilt and glycans involved in ring penetrations were removed. A 50 ns MD simulation was performed as described previously using CHARMM-GUI and GROMACS. The simulation analysis and figures generation were performed using PyMOL (v2.5) and UCSF Chimera (v1.19). The PTS3 of MUC2 chain A was modeled applying symmetry operations to the chain B PTS3 only for illustrative purposes.

## Results

### The ETEC serine protease EATA acts on the C-terminal protein region of MUC2

Previous studies have demonstrated that EatA is proteolytically active on the MUC2 mucin ^6^. To identify the region of MUC2 susceptible to cleavage we used various recombinant proteins spanning different regions of the protein. (**Fig. 1A**). EatA treatment of the N- (D1, D2, D’, D3, PTS0 and CysD1) and C-terminal (PTS3, VWCN, D4, VWC1, VWC’, VWC3, VWC4 and CK) protein regions of MUC2 and analysis by SDS-PAGE under reducing conditions indicated that EatA was only active on the protein C-terminal resulting in protein fragment of approximately 170kDa (**Fig. 1B**). The protein termini of MUC2 are responsible for dimerization via intramolecular disulfide bonds aiding to the formation of large oligomeric structures as well as internally stabilized by disulfide bonds ^17,30,31^. To determine if EatA is able to resolve dimerized MUC2, the N-terminal and C-terminal were analyzed by SDS-PAGE under non-reducing conditions. The results confirmed the resistance of the N-terminal while in contrast the C-terminal multimeric structure was degraded (**Fig. 1C**). To narrow down the region susceptible to proteolytic cleavage an additional C-terminal protein (MUC2C-NC) was tested spanning the region between the von Willebrand CN domain (VWCN) and the cysteine-knot (CK) at the C-terminal (**Fig. 1A**)^17^. Analysis by protein gel electrophoresis of the truncated C-terminal of MUC2 upon EatA treatment resulted in an unaltered band migration (**Fig. 1D**). This result indicates that the EatA cleavage site is in the N-terminal before the VWCN domain. In support of this finding an additional protein, MUC2C-N, was used. This protein was truncated at the C-terminal, containing only the N-terminus of the MUC2 C-terminal protein region of MUC2 (PTS3, VWCN, D4 and VWC1). Gel analysis revealed that only the active form of EatA resulted in a reduced band size, emphasizing that the approximate region of the cleavage site is in the PTS3-VWCN region (**Fig. 1E**). The analyses were performed under non-reducing conditions, as the EatA and MUC2C-N protein are similar in size under reducing conditions. The MUC2N and MUC2C-NC/MUC2C-N proteins lack the central mucin domain of the protein composed of multiple variable number tandem repeats (VNTR). These repeats are predominantly composed of proline, threonine, and serine (PTS) residues and are the main site of *O*-glycosylation. Protease activity was validated on two proteins composed of varying number of the PTS1 (x9) and PTS2 (x7) domain modified with either only GalNAc residues or extended to Core-1 or -2 structures with additional terminal sialylation (**Fig. 1F**) ^18^. EatA treatment of the two PTS domains did not result in degradation (**Fig. 1F**). These results indicate that EatA has proteolytic activity specifically targeted towards the MUC2 C-terminal in the region between the PTS3 and VWCN domain. Cleavage in this region results in the degradation of the oligomeric mucin structure that is normally prevented by inter- and intramolecular disulfide bonds and will resolve the mucus gel.

**Figure 1:**
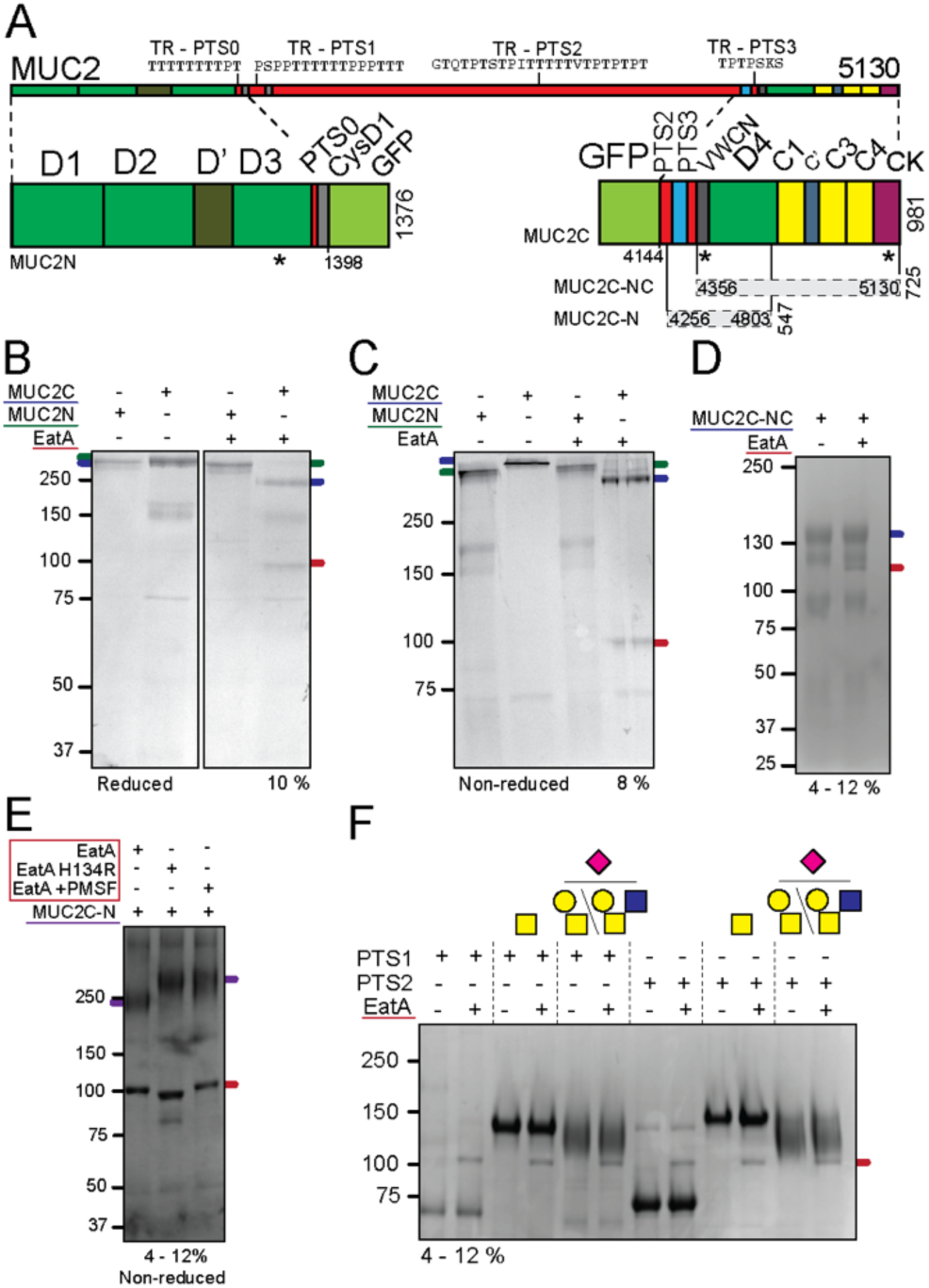
The C-terminal of MUC2 is susceptible for degradation by the protease EatA. **(A)** Schematic representation of the domain organization of the MUC2 mucin and the recombinant fusion proteins used in this study. The depicted domains are von Willebrand assembly (D), Von Willebrand C-type (C), tandem repeat mucin domain (TR-PTS), cysteine-knot (CK), CysD1 domain (CysD) and green fluorescent protein (GFP). The location of the intramolecular disulfide bonds responsible for dimerization are indicated with an asterisk (*). **(B - C)** SDS-PAGE analysis under non-reducing and reducing conditions of the recombinant MUC2N and MUC2C after EatA treatment visualized by Coomassie. **(D)** SDS-PAGE analysis and Coomassie staining of the truncated C-terminal protein MUC2C-NC treated with EatA. **(E)** MUC2C-N treated with active, non-catalytic mutant or inhibited EatA analyzed under non-reducing conditions by SDS-PAGE and visualized by Coomassie. **(F)** SDS-PAGE of the MUC2 PTS1 and PTS2 variants modified with single GalNAc residues or sialylated Core-1 or 2 oligosaccharides after treatment with the EatA protease. Color coded lines representing the position of the indicated protein on the electrophoresis gel.

### The EatA cleavage site is in the C-terminal short TR-PTS3 domain of MUC2

Proteases of the SPATE family have a broad substrate specificity predominantly towards *O*-glycosylated secreted and cell surface proteins ^32^. One of the earliest established substrates of SPATEs is IgA1 which can be cleaved in the hinge region between the Fab an Fc domain. The responsible protease is the immunoglobulin A protease (IgAP) family, which is produced by various pathogens including *Haemophiles influenzae*, *Neisseria meningitidis* and *Neisseria gonorrhoeae*. EatA is not proteolytically active on IgA1 as demonstrated by gel electrophoresis (**Fig. S1A**). The IgA1 hinge region is predominantly composed of serine and threonine residues and modified with three to six Core-1 *O*-glycans ^33^ . This region resembles the PTS repeat in MUC2 and when we performed a sequence similarity search between the IgA1 hinge region and all proteins in the human proteome MUC2 is the protein with the highest local alignment. With 73% sequence similarity with the TR-PTS3 region in the C-terminal region of MUC2 between amino acid 4329 to 4348 (**Fig.S1B**). This region of MUC2 is lacking in the truncated MUC2C-NC construct hence we did not observe any cleavage product (**Fig. 1A, D)**. To determine the exact proteolytic site of EatA in the MUC2 protein C-terminal we applied mass spectrometry to identify the neo N-terminus. The method is based on α-amine specific labeling with the TMPP reagent which retains the positive charge essential for peptide sequencing ^12,34^. The EatA treated MUC2C protein was labelled with TMPP prior to analysis by protein gel electrophoresis and stained with Coomassie. The 170 kDa band was excised and the protein was digested using trypsin. Resulting peptides were analyzed using mass spectrometry and the TMMP labelled peptide SK was identified as the neo N-terminal after EatA cleavage (**Fig. 2A)**. The identified site occurs three times in the TR-PTS3 region peptide indicating the potential for multiple cleavages (**Fig. 2B**). To confirm the cleavage site, a recombinant 34-mer peptide covering the cleavage site was synthesized corresponding to amino acid residue 4328 to 4362 and denoted as pep-PTS3. Incubation of the recombinant PTS3 peptide with EatA and subsequent mass spectrometry analysis yielded only peaks representing the multiple charged complete peptide and no cleaved product (**Fig. 2C**). As this region of the protein is likely to be glycosylated, we hypothesized that EatA could potentially be an *O*-glycoprotease dependent on the presence of *O*-glycans for activity. To determine the extent of *O*-glycosylation in this region of the protein we characterized the modified sites and *O*-glycan composition. Glycopeptides were generated from MUC2C after desialylation and digestion with the *O*-glycoprotease OpeRATOR, which acts before a Core-1 modified serine or threonine, and then analyzed by mass spectrometry. The results highlight the extend of *O*-glycosylation with 10 serines and threonines modified with either GalNAc or GalNAc-Gal (**Fig. 2D and Supp. table 1**). The occupancy is variable and only Core-1 glycans were observed due to the expression in CHO-K1 cells which are limited in their ability to synthesize extended *O*-glycan structures ^36^. This observation emphasized the extent of glycosylation around the cleavage site and its potential necessity for EatA activity. To validate this, we modified the PTS3 peptide using a combination of recombinant GalNAc-tranferases introducing between 7 and 13 GalNAcs to the 38-mer peptide. Modified sites were localized by mass spectrometry after digestion with the OgpA *O*-glycoprotease reaffirming the high site occupancy and variable rates of modification that was observed for the MUC2C protein (**Fig. 2E and Supp. table 1**) ^37^. Treatment of the GalNAc-modified peptide with EatA for up to 24 h did not results in any protease cleavage (**Fig. 2F)**. The impact of MUC2 *O*-glycosylation on protease activity was further validated by enzymatic deglycosylation of the protein backbone using sialidases, galactosidase and endo-N-Acetylgalactosaminidase sequentially or in combination. EatA treatment for 1 h was able to degrade MUC2C in presence or absence of *O*-glycans (**Fig. 2G)**. Both partial and complete removal of the glycan residue enhanced the protease kinetics, specifically the removal of the terminating sialic acid. This result is in line with the general considered role of sialic acid in protecting glycoproteins from proteases to increase protein half-life as demonstrated for other SPATEs ^38,39^. These results confirm that EatA cleaves in a region of MUC2 that is highly *O*-glycosylated, but is not dependent of glycosylation for activity.

**Figure 2:**
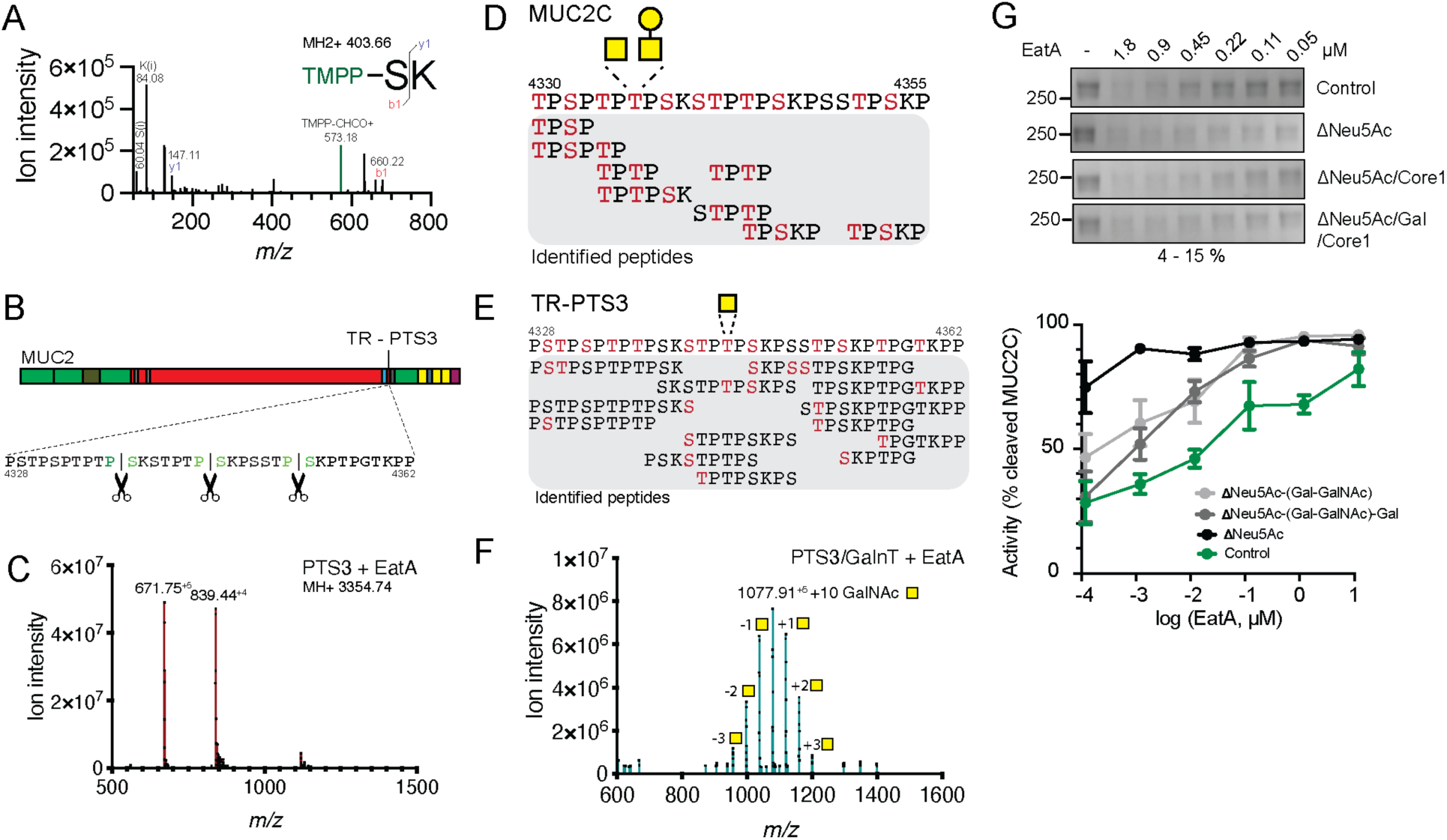
Identification of the EatA cleavage site in the MUC2 C-terminal. **(A)** Fragmentation spectra of the neo N-terminal after EatA cleavage followed by trypsin digestion identified the amino acid sequence SK. The reporter ion at 573.18 *m/z* indicates TMPP labelling, and the b-1 fragment the N-terminal serine, the peptide composition is further supported by the immonium (i) ions. **(B)** Overview of the EatA cleavage site in MUC2 and its reoccurrence in the PTS3 repeat **(C)** Mass spectra of the recombinant PTS3 spanning peptide after treatment with EatA. The MH^+4^ and MH^+5^ peaks represent the mass of the full-length 38-mer peptide at a mass of 3353.73 Da **(D)** Glycopeptides identified by mass spectrometry spanning the TR-PTS3 region in recombinant MUC2C, serines and threonines highlighted in red indicate residues that were found modified with GalNAc and GalNAc-Gal. **(E)** Glycopeptides identified by mass spectrometry of the recombinant GalNAc-tranferase modified TR-PTS3 containing peptide, serines and threonines highlighted in red indicate residues that were found modified with GalNAc **(F)** Mass spectra of the GalNAc modified recombinant TR-PTS3 peptide after treatment with EatA. The number of GalNAc residues introduced was between 7 and 13 with a majority of 10 modified sites indicated by the predominant peak at 1077.91 MH^+5^. **(G)** Kinetics of MUC2C digestion by EatA after sequential deglycosylation with sialidases, galactosidase and endo-N-acetylgalactosaminidase as compared to control, determined by SDS-page electrophoresis and represented as percentage cleaved (n=3).

### EatA activity is dependent on the structure of MUC2

The inability of EatA to degrade the recombinant peptide spanning the identified cleavage site indicates that the protease depends on the structure and folding of MUC2 for protease activity. In addition to the protease domain EatA contains additional smaller domains extending from the long beta-helix region of the passenger domain, which have been demonstrated by us and others to be essential for substrate recognition ^6,40^. The frequency and size of these domains varies between SPATE classes indicating a specific role ^41^. The largest domain with unknown function is (DUF) on the stalk region of EatA which has high structural homology with the same domain in other members of the SPATE family (**Fig. 3A**). To determine the requirement of the previously identified DUF domain for protease activity on the C-terminal of MUC2 an EatA mutant lacking the DUF domain was tested ^6^. MUC2C was incubated with decreasing concentrations of EatA or EatAΔDUF and analyzed by gel electrophoresis (**Fig. 3B**). EatA was able to degrade MUC2C even at a 1:10,000 protease to substrate ratio while EatAΔDUF was unable to degrade MUC2C, not even when using a large excess of the protease. This highlights that the DUF domain is essential for the recognition of MUC2. To determine how important the secondary structure of MUC2 is for substrate recognition we unfolded the MUC2 by reducing all disulfide bonds followed by alkylation of the free sulfhydryl groups to prevent refolding. EatA was unable to degrade MUC2C after disrupting the disulfide bonds indicating that the protease is depended on secondary protein structure in addition to the presence of a specific cleavage site (**Fig. 3C**). This is consistent with the inability of EatA to degrade the recombinant PTS3 peptide (**Fig. 2C**). The interaction between the protein and the protease was demonstrated under native conditions by size exclusion chromatography after incubation of and MUC2C-N and the catalytically impaired H134R EatA mutant for 1 h prior to separation. The partial overlap between the elution of MUC2C-N and EatA indicate that the two proteins exist as a complex (**Fig. 3D**). In contrast, when the DUF domain was lacking the interaction was not observed (**Fig. 3E**). The size exclusion chromatography results indicate that the interaction between protease and substate is transient and the enzyme concentration during the chromatography analysis. To further demonstrate the direct interaction between MUC2 and EatA, the protein complex was chemically crosslinked using glutaraldehyde and analyzed by gel electrophoresis. The crosslinking formed protein complexes far exceeding the size of the MUC2C-N dimer observed around 250 kDa indicating oligomerization between the two proteins (**Fig. 3F).** Mass spectrometry analysis of the protein composition of the four main bands observed on the gel confirmed the presence of both MUC2 and EatA in the three high molecular mass weight bands (**Fig. 3G)**. To determine the interaction between the EatA protease and MUC2 C-terminal beyond the PTS3 domain we crosslinked the mutant protease and the MUC2C-NC protein only composed of the C-terminal beyond the cleavage site (**Fig. 1A**). We observed that the combination still resulted in a reduction of the EatA band around 100 kDa and an increase in the high molecular weight bands. No crosslinking was observed in the protease lacking the DUF domain (**Fig. 3H**). This suggest that in addition to the PTS3, the DUF domain is required for protease interactions between the region beyond the cleavage site of the MUC2 C-terminal that includes the von Willebrand factor CN, D4, C1, C3 domains and cysteine knot. These results demonstrate the concerted interaction between distinct regions of both the MUC2 mucin and EatA are essential for protease activity, involving both the DUF domain on the stalk region of EatA and the domains beyond the cleavage site region of the MUC2 C-terminal.

**Figure 3:**
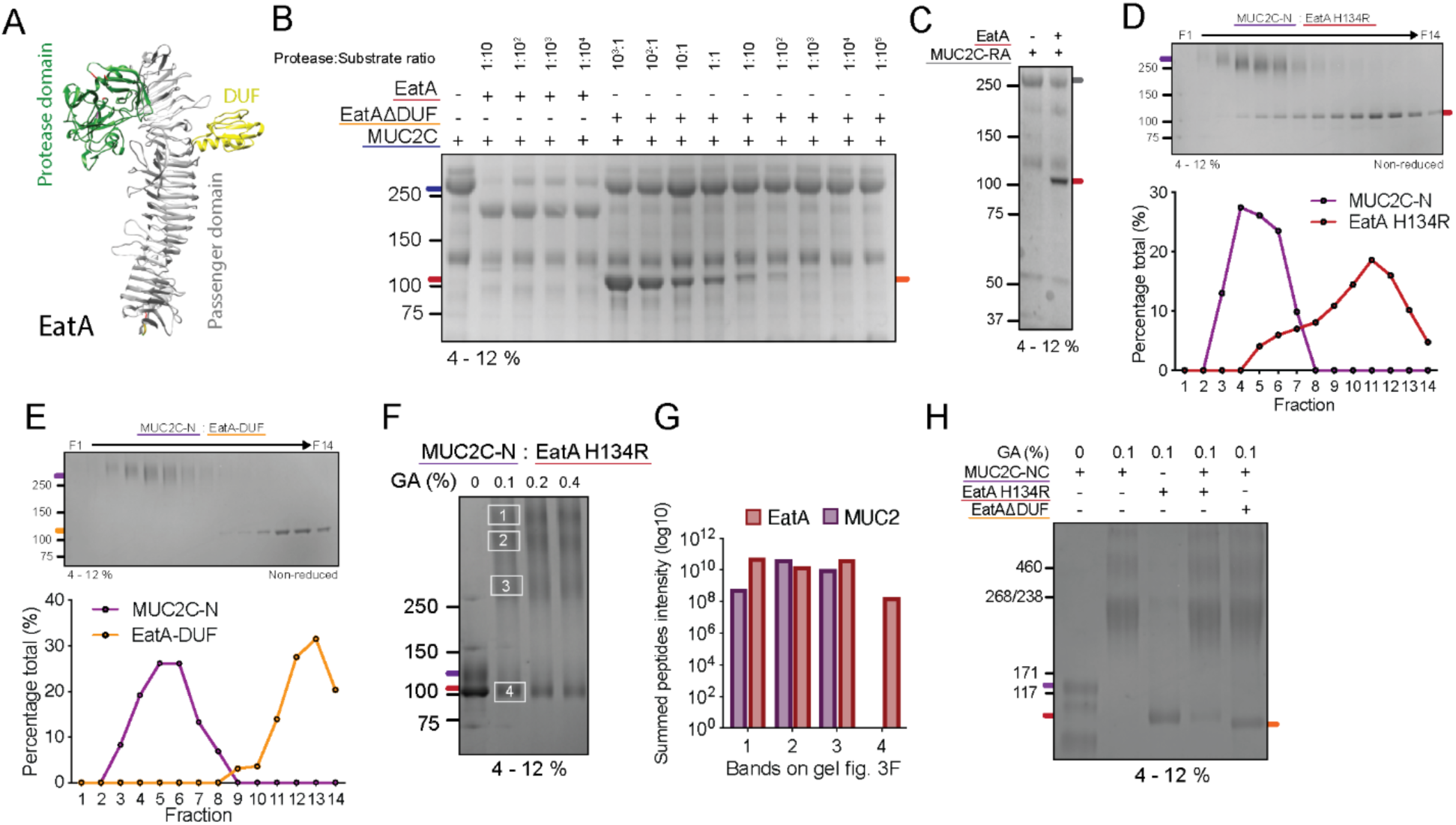
EatA substrate recognition is driven by domains and protein folding. **(A)** EatA protein structure as predicted using AlphaFold (UniProtID: Q84GK0). The serine protease domain in green, domain with unknown function (DUF) in yellow and passenger region in grey. **(B)** Protein gel electrophoresis of MUC2C treated with increasing concentrations of EatA or EatA-DUF lacking a domain extending from the helical stalk region. **(C)** SDS-PAGE analysis of the EatA protease activity on unfolded reduced and alkylated MUC2C-N (MUC2C-RA). **(D,E)** Quantification of the proportion of MUC2C-N and EatA H134R or EatAΔDUF in each fraction after 1 hr pre-incubation and subsequent size exclusion chromatography. Distribution was determined based on band intensity after protein gel electrophoresis. **(F)** Protein crosslinking of MUC2C-N and EatA H134R with increasing concentrations of glutaraldehyde (GA). **(G)** Mass spectrometry analysis of the protein components of the bands after glutaraldehyde cross-linking. Proteins were digested using trypsin and the data is represented as the summed peptide intensities for each protein in the sample. **(H)** Protein crosslinking of MUC2C-NC lacking the cleavage site region and EatA H134R or EatAΔDUF using glutaraldehyde (GA).

### EatA selectively acts on the human intestinal mucus layer

The MUC2 mucin is the major structural component of the mucus layer covering the epithelial surface in the small and large intestine. By c reating large oligomeric mesh like structures that limits the penetration of the commensal microbiota in the intestinal lumen ^42^. The pathogenesis of ETEC is depending on adherence to the epithelial cells followed by release of enterotoxins, which under normal conditions is prevented by the mucus layer. To determine if EatA is able to degrade the mucus layer to facilitate access *in vivo* we applied our *ex vivo* mucus measurement setup ^43^. Colonic tissue was mounted horizontally in a perfusion chamber kept at 37°C, the epithelium was visualized using a DNA stain and fluorescent beads in the size range of bacteria were applied apically to visualize the top of the mucus layer. The protease was added to the apical tissue surface and the thickness of the mucus layer was monitored overtime by confocal microscopy. *Ex vivo* mucus measurements of colonic tissue from wildtype mice treated with EatA for up to 20 minutes had no effect on the thickness or penetrability of the mucus layer as indicated by the constant distance between the cell surface and the beads on top of the mucus (**Fig. 4 A-B, Sup. Mov. 1**,2). These results indicate that the mucus in mice is not degraded by EatA. Since mice are not susceptible to ETEC infection we hypothesized that this is potentially due to an inability to degrade the mouse Muc2 mucin. To address the role of species specificity for EatA substrate recognition, we generated a transgenic mouse model in which we introduced the human MUC2. Human MUC2 expressing mice were crossbred with Muc2 knockout animals to generate MUC2^+^/Muc2^-/-^ mice which have a humanized mucus layer. Treatment of mucus in the MUC2^+^/Muc2^-/-^ mice with EatA resulted in a rapid degradation of the mucus layer, as shown by the decreased distance between the beads and the epithelial surface (**Fig. 4C**). Beads were observed to settle in clusters and float away in the absence of the mucus layer (**Sup. Mov. 3**). Mucus analysis without EatA addition demonstrates that the mucus of MUC2^+^/Muc2^-/-^ mice maintained its barrier function over time (**Fig. 4D, Sup. Mov. 4**). Repeated *ex vivo* analysis corroborated that only the MUC2^+^/Muc2^-/-^ mucus is susceptible for degradation by EatA (**Fig. 4E**). To confirm that the transgenic mice generate a functional mucus layer that is maintained over time we used an additional mucus measurement technique in which the thickness is measured using a penetrating needle while the mucus surface is visualized by beads ^21^. Both initial mucus thickness and growth over time were similar in C57BL/6N and MUC2^+^/Muc2^-/-^ mice, however, only mucus produced by MUC2^+^/Muc2^-/-^ mice was degraded by EatA (**Fig. 4F**). To further validate the EatA observations in the humanized mucus mouse model, we repeated the analysis using human colonic biopsies. Biopsies were obtained from control patients without a history of intestinal disease and analyzed in the same way as the mice. EatA treatment of the mucus resulted in degradation of the layer, while the mucus layer remained intact when treated with PMSF inhibited EatA (**Fig. 4G-I**, (**Sup. Mov. 5,6**)). These results demonstrated that EatA selectively degrades the human MUC2 mucin, but not mice MUC2. The cleavage site region comparison between human MUC2 and mouse Muc2 shows a clear variation in both length and amino acid composition (**Fig. 5A**). The identified cleavage site in MUC2 is present twice in Muc2 with the same residues in the P1-P1’ position (P/S) but without a lysine in the P2’ position. The lack of a positively charged residue in this position potentially impairs substrate recognition. We validated EatA activity on the mouse C-terminal of Muc2 using a recombinant protein spanning the same region of the protein as the human MUC2C-N protein (**Fig. 1A**). Western blot analysis targeting the N-terminal FLAG-tag demonstrates that only human MUC2 is degraded while mouse Muc2 remains intact (**Fig. 5B**). To determine if the mouse Muc2C-N binds to EatA we cross-linked the two proteins with increasing concentrations of glutaraldehyde. This combination resulted in a reduction of the EatA band indicating complex formation which was dependent on the presence of the DUF domain (**Fig. 5C**). In addition to humans; pigs, cows, sheep and dogs are susceptible for ETEC infections especially during neonatal development, but not mouse and rat ^44^. When comparing the PTS3 domain amino acid sequence of the effected species to that of rodents which are resistant the major variation was observed in the length of the region (**Fig 5D**). While the terminal regions are generally conserved a ∼18 amino acids insertion is present in both mouse and rat Muc2. These additional residues will affect the spatial arrangement of the cleavage site region and potential accessibility for the protease. These result shows that EatA activity is tailored to degrade mucus of its specific host species, and providing an insight into the underlying molecular pathogenesis why mice are unaffected by ETEC.

**Figure 4:**
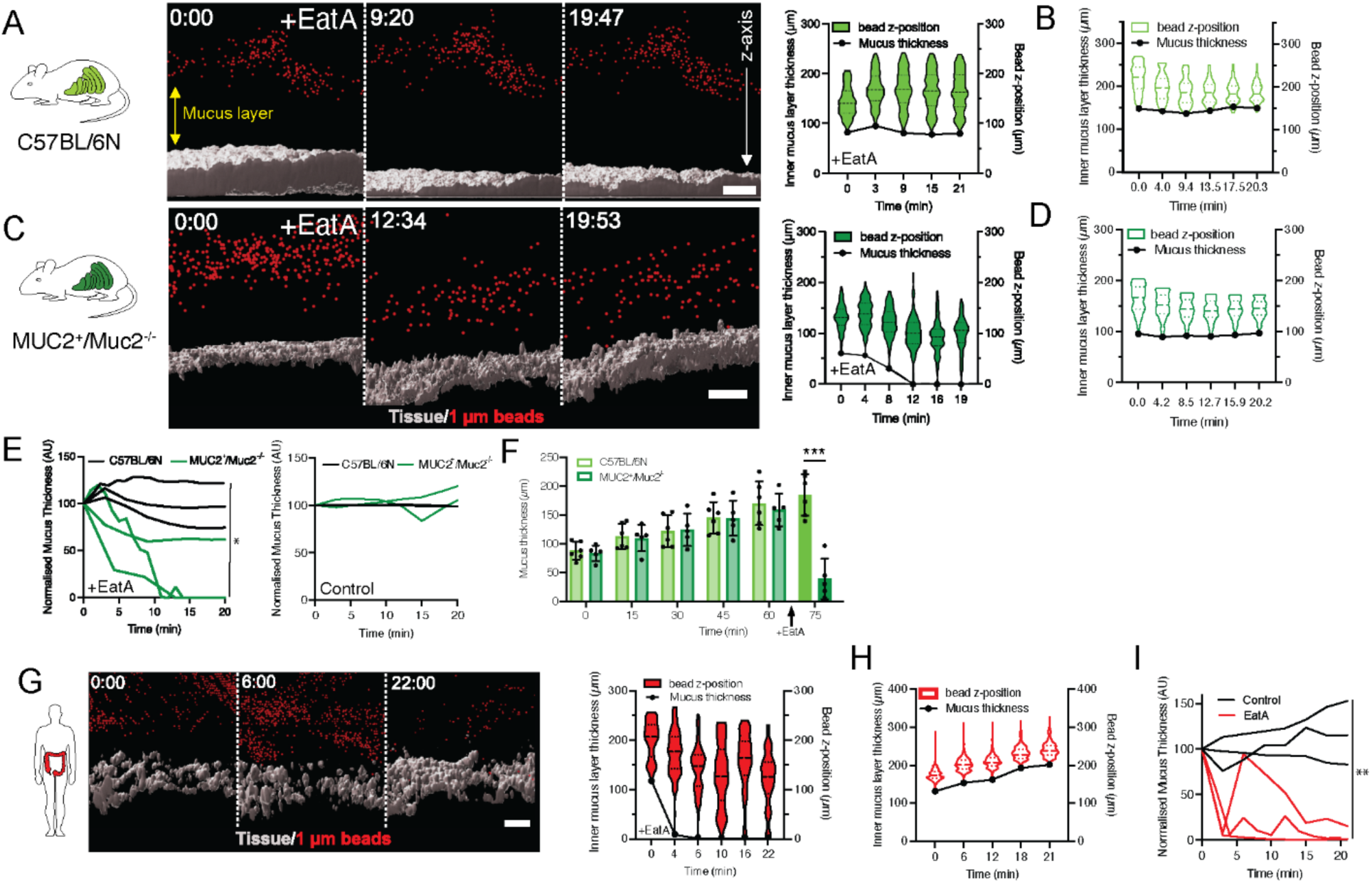
EatA specifically degrades the human colonic mucus layer. **(A)** Representative z-stack projections of *ex vi*vo mucus penetrability analysis in the colon of wildtype C57BL/6N mice upon EatA treatment over time showing the bead position (red) in relation to the tissue (grey). The violin plot shows bead frequency distribution in relation to tissue surface over time. Dashed black lines indicate the median, and solid lines the center quartiles. **(B)** Control none protease treated wildtype C57BL/6N mucus penetrability analysis over time. **(C)** Representative z-stack projections over time of ex vivo mucus penetrability analysis in the colon of MUC2^+^/Muc2^-/-^ mice upon EatA treatment. Violin plots show mucus penetrability in response to EatA treatment at each time point **(D)** Control non-protease treated MUC2^+^/Muc2^-/-^ mucus normal penetrability over time. **(E)** Combined data of replicated analysis of the penetrability assay on wildtype and MUC2^+^/Muc2^-/-^ upon EatA treatment (n=3). **(F)** Comparison of colonic mucus layer thickness and mucus growth rate over time in C57BL/6N and MUC2^+^/Muc2^-/-^ mice and the effect of EatA, error bars represent standard deviation (n= 6). **(G)** Analysis of the mucus penetrability in human colonic biopsies and the response upon EatA treatment. **(H)** and in response to PMSF inhibited EatA. **(I)** Combined data of replicated analysis of the normalized penetrability assay on patient biopsies mucus with treated with EatA or the inactivated protease (n=3). Scale bars are 50 µm. *p < 0.1 and ***p < 0.001 as determined by unpaired t-test at final timepoint E and two-way ANOVA between mouse models at each timepoint for F.

**Figure 5:**
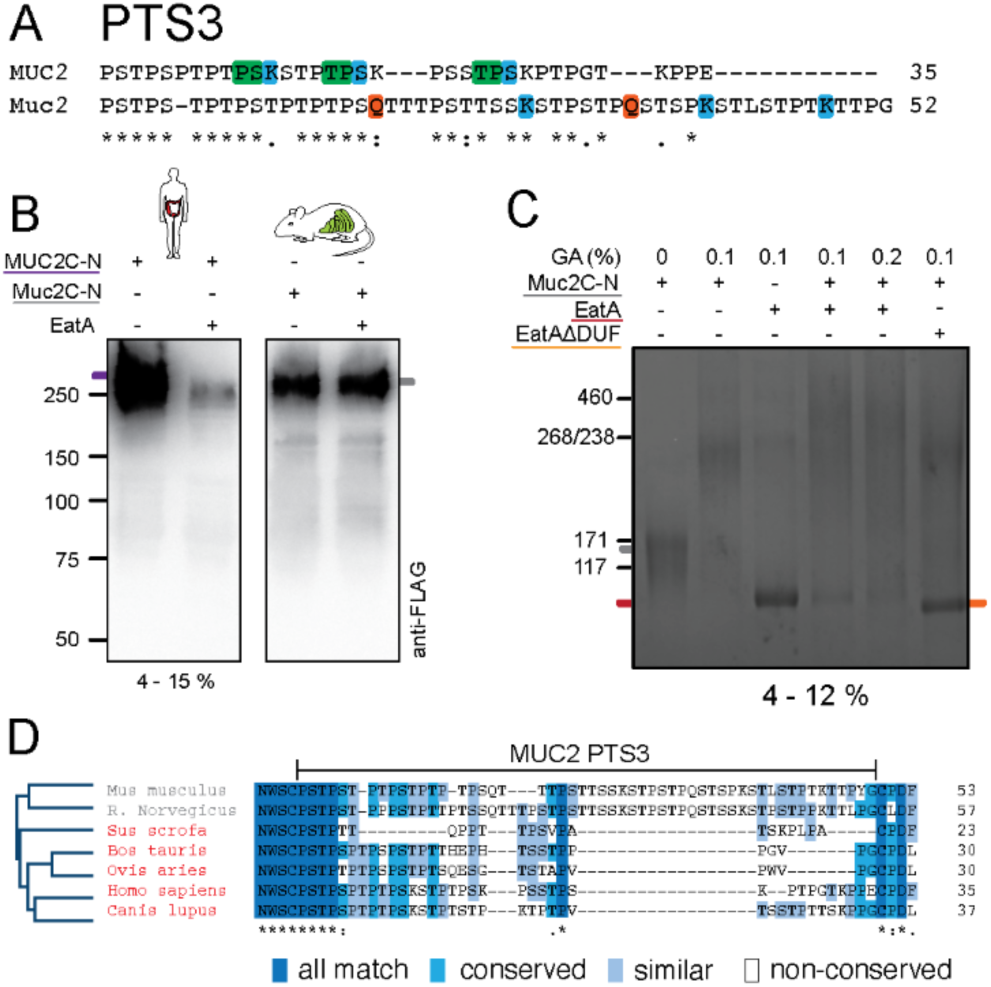
Variation in the PTS3 sequence determine species specificity. **(A)** Alignment of the PTS3 region of mouse and human MUC2. The EatA P1-P1’cleavage site position is indicated in green and lysine and glutamine in blue and red. **(B)** Western blot analysis of the human MUC2C-N and mouse Muc2C-N protein constructs treated with EatA and probed for the presence of the N-terminal Flag-tag. **(C)** Protein crosslinking of Muc2C-N and EatA with using increasing concentrations of glutaraldehyde (GA). **(D)** Phylogenetic tree and multiple sequence alignment of the MUC2 PTS3 region displaying species affected (grey) or unaffected (red) by ETEC.

### Model of the EatA binding t oMUC2

To elucidate the molecular mechanism of EatA-MUC2 recognition and cleavage, we performed docking simulations using a model based on the MUC2 VWCN-D4-VWC1 cryoEM structure (PDB: 7QCN) and a AlphaFold2 model of EatA (UniProt:Q84GK0) ^17^ . The AlphaFold2 EatA model showed very high confidence except for the DUF domain, the region suggested to drive the interaction between the two proteins. The DUF prediction presented low confidence areas in the regions connecting to the passenger domain. The predicted alignment error (PAE) between the DUF domain and the rest of the molecule was high, indicating low confidence in the relative position of the domain, typical for flexible regions. Consequently, the docking simulations were conducted using the isolated DUF domain. The highest scored complexes were analyzed extending EatA and MUC2 C-terminal (Fig. 6A) allowing small crashes that could be explained by modelling errors or flexibility and were corrected by a short (5 ns) molecular dynamics simulation. Consistently, the best scoring complexes showed the same interaction mechanism where the DUF domain was inserted in the cavity formed by the two VWCN-D4 dimers (Fig. 6B). Interestingly, the N-terminal of the VWCN from the chain A of the MUC2 monomer points towards the catalytic domain at ∼45 Å from the catalytic serine.

**Figure 6:**
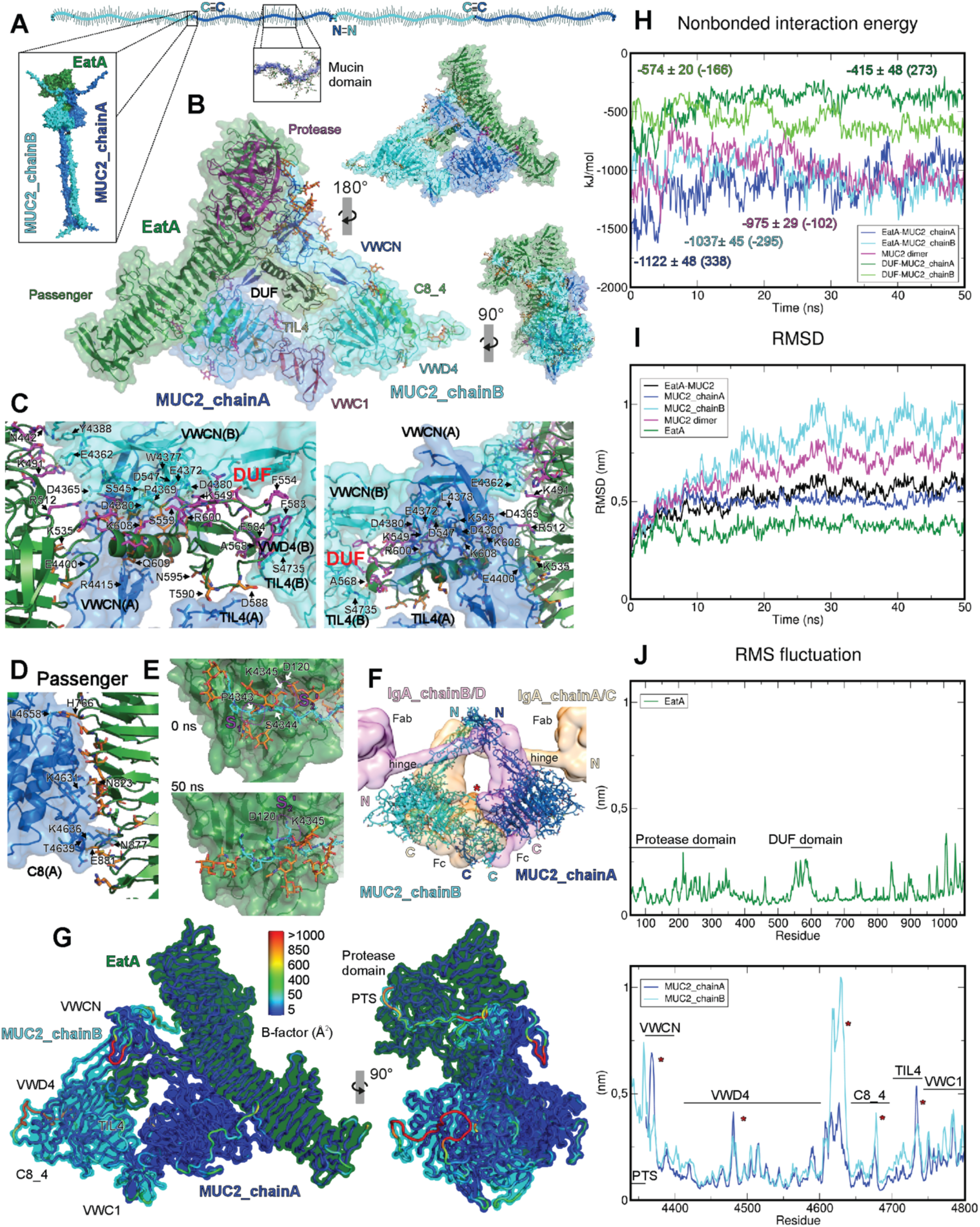
EatA-MUC2 docking model. EatA is shown in green, MUC2 chain A in blue and MUC2 chain B in cyan **(A)** Schematic sketch of a MUC2 filament drawn to scale. The *N-* and *C-* termini where the disulfide mediated oligomerization occurs are annotated. The predicted complex between the MUC2 C-terminal region (1) and EatA is showed as an inset. **(B)** EatA-MUC2 docking model after 50 ns molecular dynamics (MD) simulation, surface (excluding glycans) and cartoon. MUC2 chain A glycans are shown in magenta and chain B in orange. In the central figure cartoon representation, the different protein domains are showed in different colors: EatA Protease domain in magenta, Passenger domain in green and DUF in black; and MUC2 VWCN in blue, VWD4 in cyan, C8_4 in green, TIL4 in orange and VWC1 in red. The complex is turned clockwise by 90°on the y-axis in the bottom figure and 180° in the upper one. Both are reduced by 50% in size respect the central figure. **(C)** DUF-MUC2 and **(D)** Passenger domain-MUC2 detailed interaction. Interfacing residues are shown in detail, the most relevant ones are labeled. EatA residues interacting with MUC2 chain A are shown in orange and with chain B in magenta. Only the surface representation of MUC2 is shown. **(E)** EatA active site - MUC2 PTS4 detailed interaction before (0 ns) and after (50 ns) MD simulation. *O*-glycans are shown in orange. Only the surface representation of EatA is shown. The residues from P1 to P2’ positions are marked. S1 and S2’ pockets are annotated and D120 in S2’ is labeled. **(F)** MUC2-IgA (PDB: 1IGA) superposition. IgA chain A and C surface is shown in light orange and chain B and D in pink. MUC2 is represented by cartoons and sticks. IgA and MUC2 *N-* and *C-* termini are labeled and IgA domains are specified. The IgA region predicted to interact with IgAP is marked with a red star. **(G)** EatA-MUC2 B-factors, surface and cartoon representation. The cartoons are colored by the B-factors derived from the MD simulation as shown in the color bar. **(H)** Root mean square deviation (RMSD) over time during MD. The same figure color code is used plus EatA-MUC2 complex that is shown in black and MUC2 dimer in pink. **(I)** Nonbonded interaction energy over time during MD calculated as short-range Coulombic interaction energy plus short-range Lennard-Jones energy. The average energy value, standard deviation and global increase is shown for each curve. Same color code as **(H)**. **(J)** Root mean square (RMS) fluctuation in individual residues over the MD simulation, EatA in the top graph and MUC2 in the bottom. The different domain locations are specified. High variability regions between MUC2 chains are marked with red starts.

The PTS3 was modelled by extending it from the VWCN domain until it reached the EatA active site based on the observation that PTS domains extend linearly due to the extensive *O*-glycosylation ^45^. According to the model, the most C-terminal of the three putative cleavage sites (Pro4352-Ser4353) is too close to the VWCN domain to be cleaved. The first cleavage site (Pro4337-Ser4338) cannot be discarded, but it is the second one (Pro4344-Ser4345) that matches the distance to the active site. Therefore, the Pro4344 was manually placed in the S1 pocket and Ser4345 in S1’ (Fig. 6E 0 ns). *O*-glycans were added to all serines and threonines in PTS3 including Ser4345 in P1 position. The model was analyzed by molecular dynamics in order to evaluate the docking prediction, determine the stability of the complex, and to improve the model. The molecular dynamics simulation disclosed that the interaction between the catalytic domain and the PTS3 is mainly stabilized by Lys4346 in P2’ forming a salt bridge with Asp120 in S2’ (Fig 6E 50 ns). The interactions involving P3-P1 positions vanished during the simulation and the S1 pocket became less pronounced. Obvious interactions between glycans and the protease domain are not observed, but neither major steric impediments.

The EatA-MUC2 model only suffered small rearrangements during the simulation, the interaction surfaces involving the DUF, passenger and protease domains were preserved (Fig. 6B). As highlighted before, the main interaction occurs between EatA DUF domain and MUC2 chain A VWCN(A) and TIL4(A) domains; and MUC2 chain B VWCN(B), VWD4(B) and TIL4(B) domains (Fig. 6C). This region of MUC2 is not protected by *N*- or *O*-glycans that could affect the dimer formation and presents a hydrophobic region between the VWCN(B) and the VWD(B) domains where the hydrophobic residues from tip of the DUF domain (Phe554, Phe583 and Phe584) are predicted to be positioned. The interaction is further stabilized by multiple hydrogen bonds and salt bridges along the surface of the DUF domain and the adjacent regions of the passenger domain, the VWCN domains from both MUC2 monomers and the TIL4 domain from chain B. Approximately 1500 Å^2^ of surface area is buried in this interaction. Remarkably, the MUC2-EatA interacting region highly resembles the IgA1 interface predicted to interact with IgAPs SPATEs of *Haemophilus influenzae* and *Neisseria gonorrhoeae* (Fig. 6F) ^46^. The model reveals a secondary interaction surface that spans around 600 Å^2^ between MUC2 C8 (A) and the EatA passenger domain central region among the DUF domain and the C-terminal (Fig. 6D). The interaction is purely hydrophilic and it could have a role in substrate recognition even though the model suggest that the contacts are weaker and much less stable than established by the DUF domain.

The current model thus supports that EatA recognizes a lysine in position P2’ and cleaves the PTS3 not closer than ∼15 amino acids N-terminally of the VWCN domain in an *O*-glycosylation-independent manner. The analysis of the nonbonded interaction energy (short-range Coulombic interaction energy and short-range Lennard-Jones energy) during the molecular dynamics simulations confirmed that the complex reached an equilibrium after ∼15 ns (Fig 6H). The EatA-MUC2 chain A interaction energy got worse during the initial steps of the simulation but was compensated by the improvement in the EatA-MUC2 chain B interaction energy, so globally the nonbonded energy remained stable. The root mean square deviation (RMSD) evolution over time also reached an equilibrium between 15-20 ns (Fig 6I). The RMS fluctuation (RMSF) in individual residues (Fig 6G,J) explains the variances observed in the RMSD evolution in the different chains over the time. EatA shows low RMSF except for a loop in the protease domain, the flexible DUF domain and the C-terminal region, resulting in a low average RMSD. On the other hand, the flexible regions in MUC2 showed much higher RMSF increasing the overall RMSD. The effect is more evident in the chain B where the PTS domain and the connecting loop between the VWD4 and C8_4 domains present very high mobility. The comparison between MUC2 chains reveals some of the regions interacting with EatA. They show less flexibility compared to the other chain suggesting that the interactions are stable. In MUC2 chain B, the N-terminal region of VWCN, the TIL4 domain and a region in the VWD4 shows reduced RMSF due to the interaction with the EatA DUF domain. In contrast, the chain A is mainly stabilized by the interaction with the passenger domain in the C8_4 domain and the connecting loop between the VWD4 and C8_4 domains. The model highlights the importance of the spatial arrangement between the DUF and the protease domain in engaging the cleavage site region of MUC2.

## Discussion

Enteric pathogens need to overcome the largely impenetrable mucus layer to engage with the intestinal epithelium^47^ . Here we demonstrate that ETEC can resolve the mucus layer utilizing the highly specialized SPATE protease EatA that directly acts on the main protein component of intestinal mucus, the MUC2 mucin. MUC2 forms oligomers that are highly resistant to proteolysis in the hostile environment of the intestinal lumen, leaving only a few regions susceptible to protease cleavage, which would result in disruption of the mucus gel ^11,12^ . The EatA cleavage site is localized to a mucin domain in the C-terminal part of the protein closely resembling the central mucin domain, which has been considered to be proteolytically resistant due to its extensive *O*-glycosylation. In contrast to other recently identified *O*-glycoproteases acting on mucin domains EatA is not dependent on glycosylation but tolerates glycans in proximity which was supported by molecular dynamics analysis ^48–50^. The recombinant proteins used in this study were produced using CHO-K1 cells expressing only a subset of glycosyltransferases limiting the glycosylation to sialylated Core-1 structures ^12^. *O*-glycan structures identified on MUC2 in the human intestine are predominantly based on Core-3 and far more branched and extended ^51^. Our *ex vivo* mucus analysis on human mucus highlight that even the core structure or the potential proximity of extended *O*-glycans structure does not prevent EatA activity. *O*-glycans on mucin domains are dense and heterogenous which hinders efficient recognition by proteases. EatA is unable to act solely on the synthetic peptide covering the cleavage site in the PTS3 region itself suggesting additional binding sites are essential for stabilizing the substrate-enzyme complex. Previously one putative binding domain (DUF) was identified to be essential for MUC2 recognition and degradation ^6^ which we here confirm binds with high specificity to a cleft C-terminally of the PTS3 in between the two VWD4 domains of the MUC2 homodimer. In the absence of the DUF, no cleavage activity was observed and binding to MUC2 was reduced highlighting the importance of the DUF domain for initiating interactions with its substrate. DUF domains are present in many members of the SPATEs family and the domain is the likely key determinant of substrate specificity ^52^. Earlier assumptions of species tropism of enteric pathogens is suggested to be dependent on the presence of the receptors for the enterotoxin and host specific adhesins ^44^. The lack of EatA activity on mouse mucus indicates that species variation in the MUC2 sequence also determines host selection. Mucus is one of the barriers ETEC will require to overcome during infection and could therefore be considered a major contributor of host selection. Despite DUF dependent binding to mouse Muc2 no cleavage occurs, supporting the concept that variation in the sequence of the PTS3 region determines specificity. The lack of cleavage site and variable length in the PTS regions of mouse Muc2 are indicators as to why mice are not a suitable model for pathogenic *E. coli* infection ^53^. In contrast, *Citrobacter rodentium* commonly used as a mouse model for Enteropathogenic *E. coli,* expresses Pic a SPATE involved in colonization ^54^. *Citrobacter rodentium* infection of Muc2^-/-^ mice is lethal underscoring the crucial role of the mucus layer ^55^. Pic is similar in domain organization to EatA with a slight variation in the active site and spatial arrangement of the domains along the autotransporter domain to facilitate selective Muc2 cleavage. The high level of substrate specificity among SPATEs suggests that previous studies relying on crude mucus extractions from non-host species provide limited insights into the actual substrate, the mode of interaction and its function *in vivo*. To overcome this limitation, we generated a transgenic model selectively expressing human MUC2 which have a normal functional mucus layer under baseline conditions. EatA treatment efficiently removed the mucus from chimeric mice at a similar rate to that seen in human colonic biopsies, we choose to perform this analysis on intact colonic tissue for the purpose of visualization ^21^ . The colonic mucus layer is similar in protein composition and *O*-glycosylation as the small intestine and the increased thickness allows improved visualization of mucus degradation ^56^ . This novel mouse model has the potential to systematically evaluate the contribution of the microbial mucus degrading proteases during infection of not only ETEC but other human intestinal pathogens. The necessity to develop a strategy to overcome the mucus layer suggests that many SPATEs are tailored to degrade MUC2 including SepA from *Shigella flexerni* and Pic from enteroaggregative *E. coli* ^5,57^. Identifying a shared mechanism between *E. coli* pathotypes and related microbial genera has the potential for the development of vaccine strategies to block the initial binding to MUC2 thereby preventing mucus degradation and colonization ^8^. Identification of the cleavage site and substrate recognition is essential for the progress of such potential developments.

In conclusion, we present a mechanistic insight into the degradation of MUC2 by the ETEC protease EatA. Localizing the cleavage site to a susceptible region of MUC2 that results in mucus gel solubilization, and how variation in the cleavage site region between species drives host selectivity. Protease activity in the unstructured region of MUC2 is depending on the initial binding to a von Willebrand domain downstream of the cleavage site to stabilize and direct the flexible region to the catalytic site. The combined results underscore the necessity for highly specialized proteases to degrade mucus, limited in expression to intestinal pathogens and offering novel therapeutic strategy for the prevention of ETEC infections.

## Supporting information

Sup. Mov. 2

Sup. Mov. 1

Sup. Mov. 4

Sup. Mov. 3

## **A**cknowledgements

This work was supported by the Swedish Research Council grant 2020-02536 (SvdP) 2023-02474 (G.C.H), Jeansson Foundations (S.vd.P), Swedish Society for Medical Research (Svenska Sällskapet för Medicinsk Forskning) (S.vd.P), Knut and Alice Wallenberg Foundation (2017.0028) (G.C.H.), Magnus Bergvall Foundation (S.vd.P), Wilhelm and Martina Lundgren’s Foundation 2023-SA-4357 (S.vd.P), European Research Council (ERC) (101100663, 694181) (G.C.H), IngaBritt and Arne Lundberg Foundation (2028-0117) (G.C.H), Sahlgren’s University Hospital (ALFGBG-440741, ALF agreement 236501) (G.C.H). We acknowledge the Mammalian Protein expression Core Facility for recombinant protein expression and are indebted to the colonoscopists and nurses at GEA of the Sahlgrenska University Hospital.

## Author contributions

B.D., S.T-M., G.C.H., and S.v.d.P. conceived and designed the study. J.K.G., L.A., T.J.V, H.H.W, J.M.F., contributed with experimental data and interpretation. B.D., S.T-M., and S.v.d.P. wrote the manuscript, and all authors edited and approved the final version.

## Correspondence and requests for materials

Should be addressed to Sjoerd van der Post

## Data availability

The mass spectrometry proteomics data have been deposited to the ProteomeXchange Consortium (http://proteomecentral.proteomexchange.org) via the PRIDE partner repository with the dataset identifier PXDXXXX.

---

**Figure S1:**
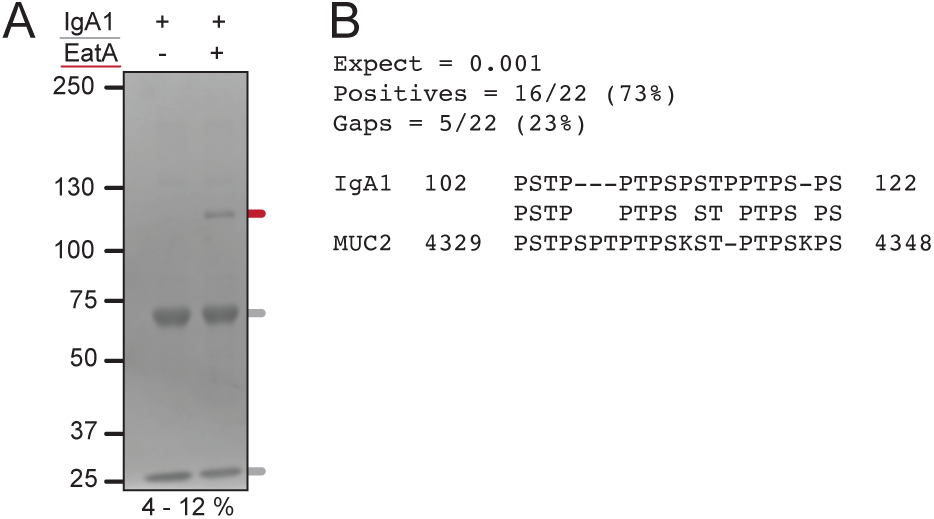
The EatA protease is active in a region similar in sequence as the hinge region of IgA1. **(A)** Protein gel electrophoresis of IgA1 treated with EatA detected with Coomassie. The heavy and light band of IgA1 are marked with in a gray line and EatA red. **(B)** Local alignment results performed using BLASTP demonstrating the sequence similarity between the hinge region of IgA1 and the TR-PTS3 in the C-terminal of MUC2.

**Figure S2:**
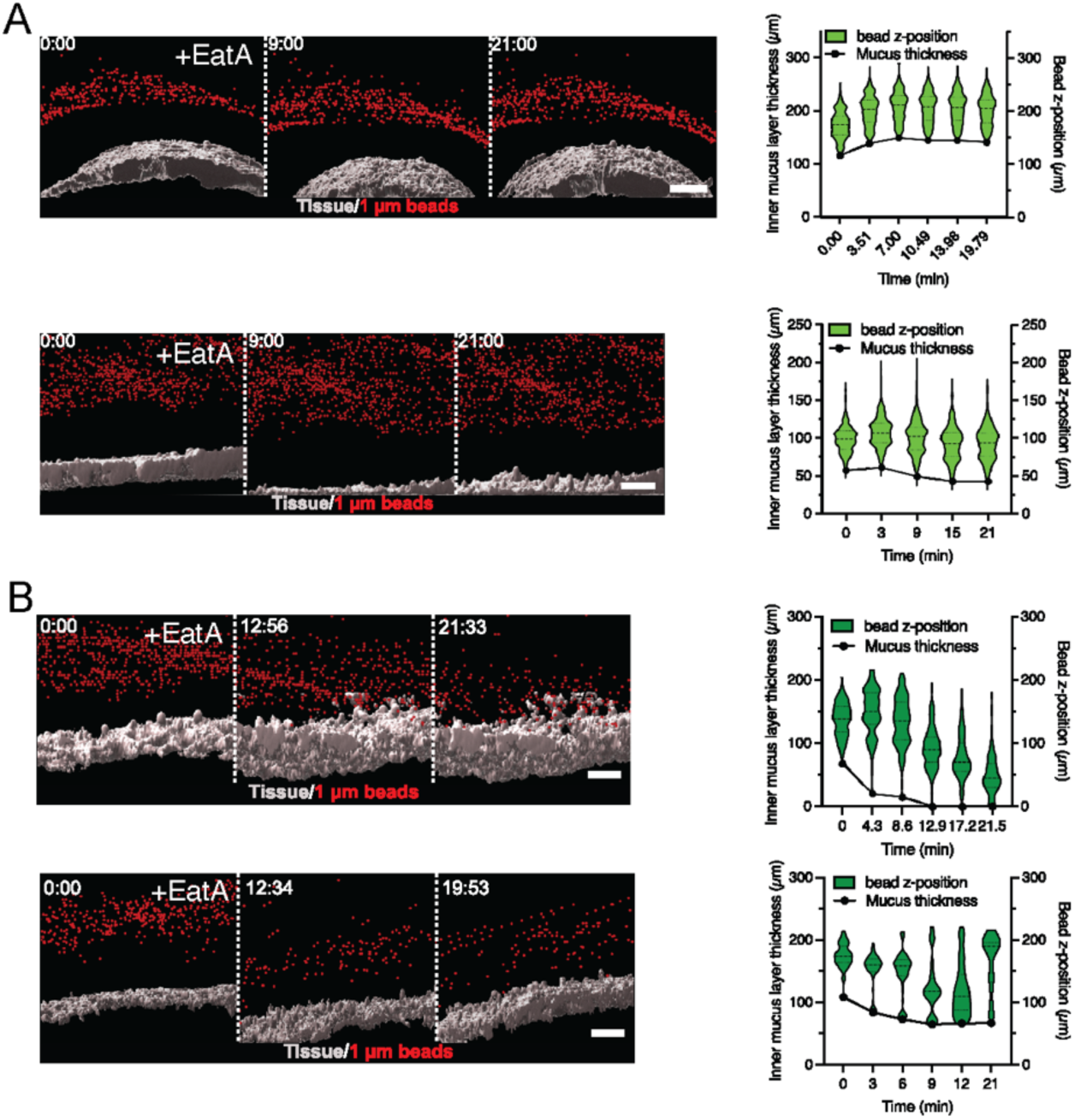
Transgenic MUC2 expressing mice produce a functional colonic mucus layer susceptible to degradation by the ETEC protease EatA. **(A)** Replicate analysis of the mucus penetrability assay performed on mouse control C57BL/6N mucus treated with EatA for 20 minutes, summarized in figure for 4D (n=3) **(B)** Replicate analysis of the mucus penetrability assay performed on mouse MUC2/Muc2^-/-^ mucus treated with EatA, summarized in figure for 4D (n=3). The violin plot show bead frequency distribution in relation to tissue over time. Dashed black lines indicate the median, and solid lines the center quartiles. Scale bars are 50µm.

